# Analysis of beta-lactam heteroresistance in CRE suggests a stage in the spectrum of antibiotic resistance

**DOI:** 10.1101/2020.09.19.304873

**Authors:** Victor I. Band, David S. Weiss

## Abstract

Antibiotic resistance is a growing crisis that threatens many aspects of modern healthcare. Dogma is that resistance often develops due to acquisition of a resistance gene or mutation, and that when this occurs, all the cells in the bacterial population are phenotypically resistant which we term “homogenous resistance”. In contrast, heteroresistance (HR) is a form of antibiotic resistance where only a subset of cells within a bacterial population are resistant to a given drug. These resistant cells can rapidly replicate in the presence of the antibiotic and cause treatment failures. If and how HR and homogenous resistance are related is unclear. Using carbapenem-resistant Enterobacterales (CRE), we show that HR to beta-lactams develops over years of antibiotic usage and that it is gradually supplanted by homogenous resistance. This suggests the possibility that HR may often develop before homogenous resistance and frequently be a stage in its progression, representing a major shift in our understanding of the evolution of antibiotic resistance.

## Introduction

Heteroresistance (HR) is a poorly understood and underappreciated form of antibiotic resistance, wherein a bacterial strain contains a resistant subpopulation within a larger susceptible population [1]. This phenotypic form of resistance is distinct from conventional “homogenous” resistance in which all bacterial cells in a population display resistance to an antibiotic. Further, the resistant cells in a heteroresistant population can rapidly replicate in the presence of a given antibiotic, in contrast to persister cells which are a distinct type of resistant subpopulation that is quiescent and cannot rapidly replicate when cultured with the drug [2]. Finally, while the resistant population in HR is enriched upon growth in a given antibiotic, the resistant cells reversibly return to baseline frequencies after subsequent drug-free subculture, indicating that the resistant subpopulation are not stable mutants.

HR has been described to many antibiotics and among many bacterial species [1,3–5], though the *in vivo* relevance of the phenomenon has been unclear [6]. However, recent findings have demonstrated that HR can mediate *in vivo* treatment failure [7–9] even though the resistant subpopulation of cells makes up a minority of the total population. Additionally, HR isolates harboring a very low frequency of resistant cells (<1 in 10,000) are usually misclassified as antibiotic susceptible, which could lead to inappropriate therapy. The origins of HR are unclear, as is the relationship between HR and conventional “homogenous” resistance in which all the cells in a population are phenotypically resistant.

To gain insight into the relationship between HR and homogenous resistance, we assessed the proportion of isolates exhibiting these phenotypes to beta-lactam antibiotics among a pool of carbapenem-resistant Enterobacterales (CRE) from Georgia, USA area hospitals. This pool of isolates was collected from 2013-2015 as a part of the Multi-site Gram-negative Surveillance Initiative (MuGSI), with clear inclusion criteria [10] to ensure a representative sampling of hospital isolates.

## Results

We used the population analysis profiling (PAP) method to assess the presence of HR and detect resistant subpopulations of cells [9]. The PAP method consists of plating bacteria on solid media on a range of concentrations of antibiotic, and assessing proportions of the bacterial population growing on each dose of drug. We tested 104 CRE from the Georgia MuGSI collection against 10 beta-lactam antibiotics which were first used clinically in the years ranging from 1961 (ampicillin) to 2015 (ceftazidime-avibactam). To minimize the chance of clinical samples containing two distinct bacterial isolates (mixed cultures), we tested each isolate after growth from a single colony. The resulting PAP curves for each isolate and antibiotic were classified as susceptible, resistant (homogenous resistance) or heteroresistant using previously determined criteria [8] and as described in the Materials and Methods. Unsurprisingly, rates of susceptibility were highest for drugs with fewer years of clinical use, while the highest rates of homogenous resistance were observed for drugs with the most years of clinical use (Figure 1A). Accordingly, there was a significant negative correlation between the years since clinical introduction of an antibiotic and the proportion of susceptible isolates (Figure 1B), and a significant positive correlation with the proportion of resistant isolates (Figure 1C). In contrast, the proportion of isolates heteroresistant to an antibiotic was highest for the drugs with an intermediate number of years since clinical introduction; over 50% of isolates were HR to meropenem, imipenem and cefepime which have been in clinical use for 10-22 years (Figure 1A). This surprising trend suggests that after introduction of a new drug, there is an initial increase in the proportion of isolates exhibiting HR, a peak, and then a subsequent decrease, as HR is supplanted by homogenous resistance. Overall there was a significant negative correlation between the proportion of heteroresistant isolates and the years since clinical introduction of the drug (Figure 1D).

**Figure 1.**
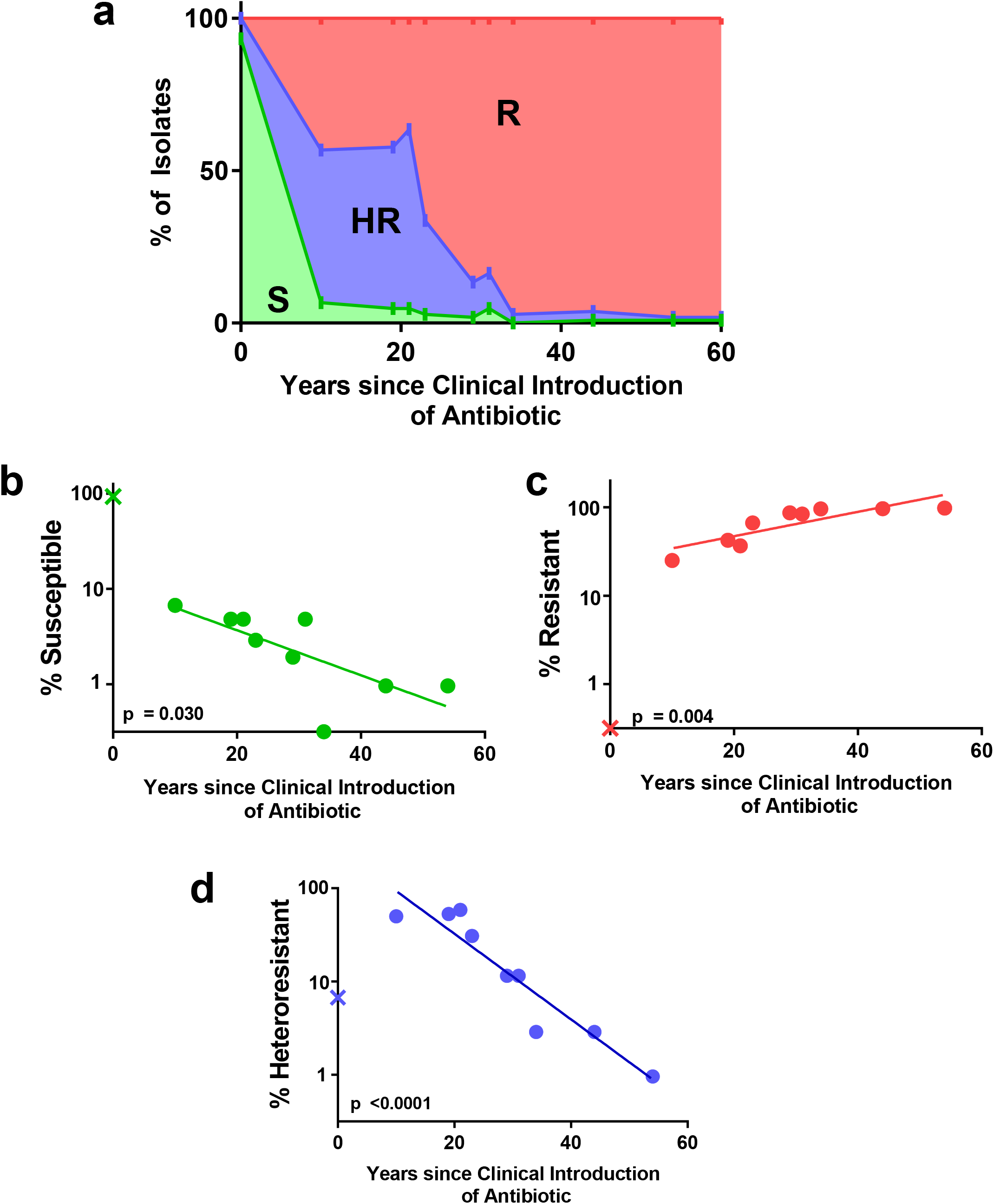
Incidence of heteroresistance and resistance correlates with number of years of clinical use of antibiotics. **a**, 104 isolates of CRE from Georgia area hospitals were assayed for heteroresistance, resistance, and susceptibility to 10 beta-lactam antibiotics which were introduced clinically in different years (Ampicillin, 1961; Cefazolin, 1971; Amoxicillin/Clavulanate, 1981; Ceftazidime, 1984; Aztreonam, 1986; Piperacillin/Tazobactam, 1992; Cefepime, 1994; Meropenem, 1996; Doripenem, 2005; Ceftazidime/Avibactam, 2015). Percent incidence of each designation was plotted by the age of the drug. Green area, Susceptible (S); Red area, Resistant (R); Blue area, Heteroresistant (HR). **b-d**, Incidence of (**b**) susceptible, (**c**) resistant, and (**d**) heteroresistant isolates by clinical age of antibiotic. Linear regression shown with p value for slope significance. Ceftazidime/Avibactam marked with ‘X’ was introduced after the study isolates were collected and was not included in linear regression analysis.

When the Enterobacterales isolates were separated by genus, we observed similar patterns for each, where *Escherichia, Enterobacter* and *Klebsiella* all displayed a significant negative correlation between incidence of HR and years of clinical antibiotic use (Figure 2A). A similar positive correlation was observed for all genera between incidence of resistance and years of clinical antibiotic use (Figure 2B). These data indicate that the HR phenomenon described here is not genus specific but rather that it occurs across all of the different Enterobacterales represented in this study.

**Figure 2.**
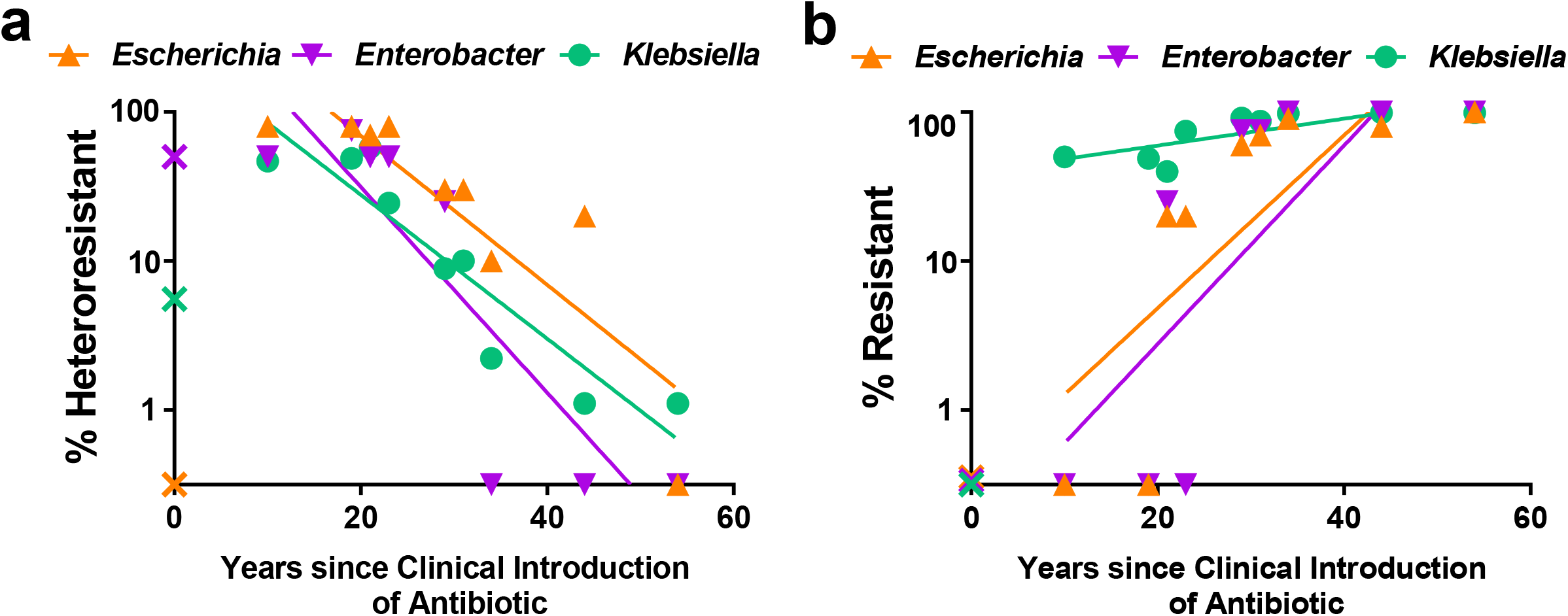
Incidence of resistance and heteroresistance by genus. **a-b**, 104 isolates of CRE from Georgia area hospitals were assayed for heteroresistance and resistance to 10 beta-lactam antibiotics and separated by genus: *Escherichia* (n=10), *Enterobacter* (n=4), and *Klebsiella* (n=90). Percent incidence of heteroresistance (**a**) and resistance (**b**) among the 3 genera is shown. Linear regressions are shown with p values for slope significance: (**a**, *Escherichia* = 0.0023, *Enterobacter* = 0.0047, *Klebsiella* = 0.0003; **b**, *Escherichia* = 0.0162, *Enterobacter* = 0.0204, *Klebsiella* = 0.0127). Ceftazidime/Avibactam marked with ‘X’ was introduced after the study isolates were collected and was not included in linear regression analysis.

## Discussion

Here, we used an incidence- and population-based approach to study resistance to 10 beta-lactams across clinical isolates of CRE from Georgia, USA. These isolates allow us to look at a snapshot of the collective resistance profile during the study years of 2013-2015. While we cannot track the development of resistance in individual isolates, we can observe proportions of resistance and consider the years of clinical use of drugs to estimate total antibiotic exposure. These results showed that for drugs in clinical use for the shortest amount of time, susceptibility dominated, while resistance dominated for those with the longest clinical use. Importantly, HR was most common to drugs introduced in the intermediate years, suggesting its role as a possible intermediate step in the development of resistance. In the future, our incidence-based study would be complemented by longitudinal studies which follow related isolates over time to observe the development of resistance in sets of related isolates. Additionally, confirmation of the role of HR in the progression of antibiotic resistance, and distinction between these pathways, will need to be explored experimentally with *in vitro* evolution experiments, which may prove challenging due to the timescales observed in this study occurring over many years.

These data have significant implications for our understanding of the development of antibiotic resistance, suggesting that HR could often be an evolutionary stage in the development of homogenous resistance (Figure 3). This model would represent a major shift from the dogmatic pathway to resistance, in which a new resistance gene or accumulation of mutations is thought to be exhibited by all the cells in the population [11]. There are at least two obvious pathways to homogenous resistance that would involve HR either directly or indirectly. The first is that HR could be a direct stage in the progression to homogenous resistance, where the genetic changes causing HR are built upon, and after acquisition of further genetic changes, 100% of the cells in the population exhibit resistance. In such a scenario, reversal of the initial genetic changes causing HR would likely result in some of the cells in the resistant strain becoming susceptible, and thus the strain would appear HR once again. The second pathway is an indirect role for HR, where HR is an intermediate step that facilitates survival of sufficient cells in the total population which then acquire distinct genetic changes that cause homogenous resistance. In this model, the genetic changes causing HR play a supportive role, and once homogenous resistance is achieved, their reversal would not have an impact on resistance. These direct and indirect roles for HR in the progression to homogenous resistance could each occur in different isolates. It is important to note that this study focuses only on beta-lactams, and that the observed trends may not apply to other antibiotic classes and warrants further study. If HR is indeed often an initial evolutionary step towards homogenous resistance, it may then represent an important source of antibiotic resistance among clinical isolates.

**Figure 3.**
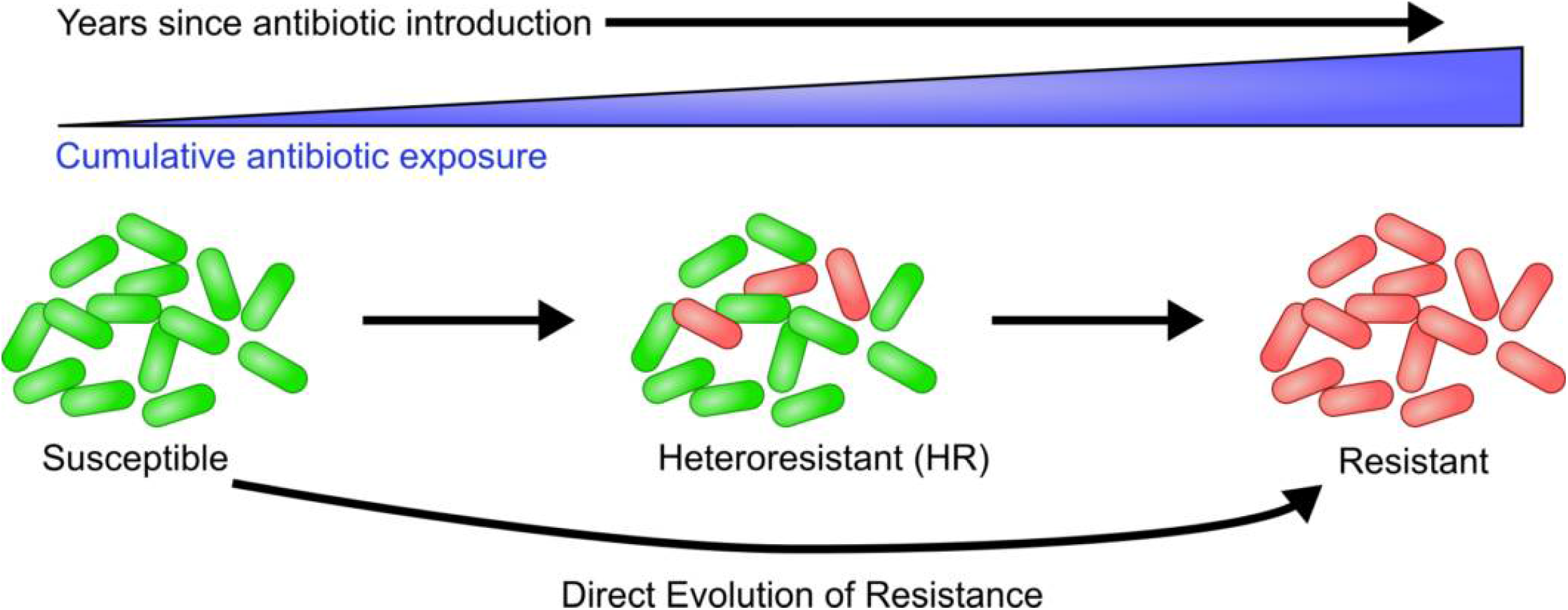
Proposed model for the evolution of resistance through heteroresistance. With increasing years after the introduction of an antibiotic, bacterial strains are cumulatively exposed to an increasing amount of the antibiotic. Strains are initially susceptible (green cells), but antibiotic exposure leads to the development of a resistant subpopulation (red cells). Increased antibiotic exposure eventually leads to homogenous resistance. Direct evolution of resistant isolates from susceptible isolates is also indicated.

Previous studies have demonstrated that roughly half of HR involves a low proportion of resistant cells and is generally undetected by current clinical susceptibility testing methods [7, 9], suggesting that the initial steps foretelling the development of homogenous resistance may be missed. Supporting this concept is our finding that for the most recently introduced antibiotic in the study, ceftazidime/avibactam (introduced in 2015), we observed several heteroresistant isolates with low frequency resistant subpopulations but no isolates exhibiting homogenous resistance. Therefore, low frequency and clinically undetected HR may be the initial harbinger of impending resistance that could be surveyed epidemiologically to inform public health and antibiotic prescribing practices.

## Materials and Methods

### Sample Collection

The 104 isolates used in this study were collected by the Georgia Emerging Infections program (GA EIP), as a part of the Centers for Disease Control (CDC) Multi-site Gram Negative Surveillance Initiative (MuGSI), detailed in Guh et al [10]. A CRE case was defined as a carbapenem-nonsusceptible and extended-spectrum cephalosporin-resistant (ceftriaxone, ceftazidime, ceftizoxime, and cefotaxime) *Escherichia coli, Klebsiella aerogenes, Enterobacter cloacae* complex, *K pneumoniae*, or *K. oxytoca* isolate recovered from a body site that is normally sterile or urine from individuals residing in the surveillance area during January 2013-December 2015. Isolates were collected by participating Georgia hospitals, and tested for inclusion criteria by GA EIP and CDC.

### Population Analysis Profile

Population analysis profile was performed as previously described [9]. Isolates were grown up from single colonies overnight and plated on 6 plates containing 0, 0.25, 0.5, 1, 2 or 4 fold the breakpoint amount of antibiotic. Growth from single colonies minimizes the chances of assessing mixed cultures of bacteria that can be found in clinical isolates. The breakpoints used were defined by CLSI [12] as the minimum MIC for resistant isolates. Colonies were enumerated and survival was calculated by comparing to a drug free plate. Isolates were classified as previously described [9] and as indicated in Figure S1; susceptible if they did not survive at 1x the breakpoint, resistant if more than 50% survived at or above 1x the breakpoint, and heteroresistant if at least 1 in 10^-6^ but fewer than 50% of cells survived at 2x the breakpoint or above. Setting the threshold for HR at 1 in 10^-6^ makes the inadvertent inclusion of stable resistant mutants unlikely, since they would need to be present at a high level representing an extremely high mutation rate. Each isolate was tested against 10 beta-lactam antibiotics: ampicillin, cefazolin, amoxicillin/clavulanate, ceftazidime, aztreonam, piperacillin/tazobactam, cefepime, meropenem, doripenem, ceftazidime/avibactam. Raw CFU counts and interpretations are available in Extended Data S1.

### Data Analysis

Year of clinical introduction for each antibiotic used in this study was determined by the approval date by the US Food and Drug Administration. Correlations with year of clinical introduction were calculated using a linear regression analysis.

---

**Figure S1.**
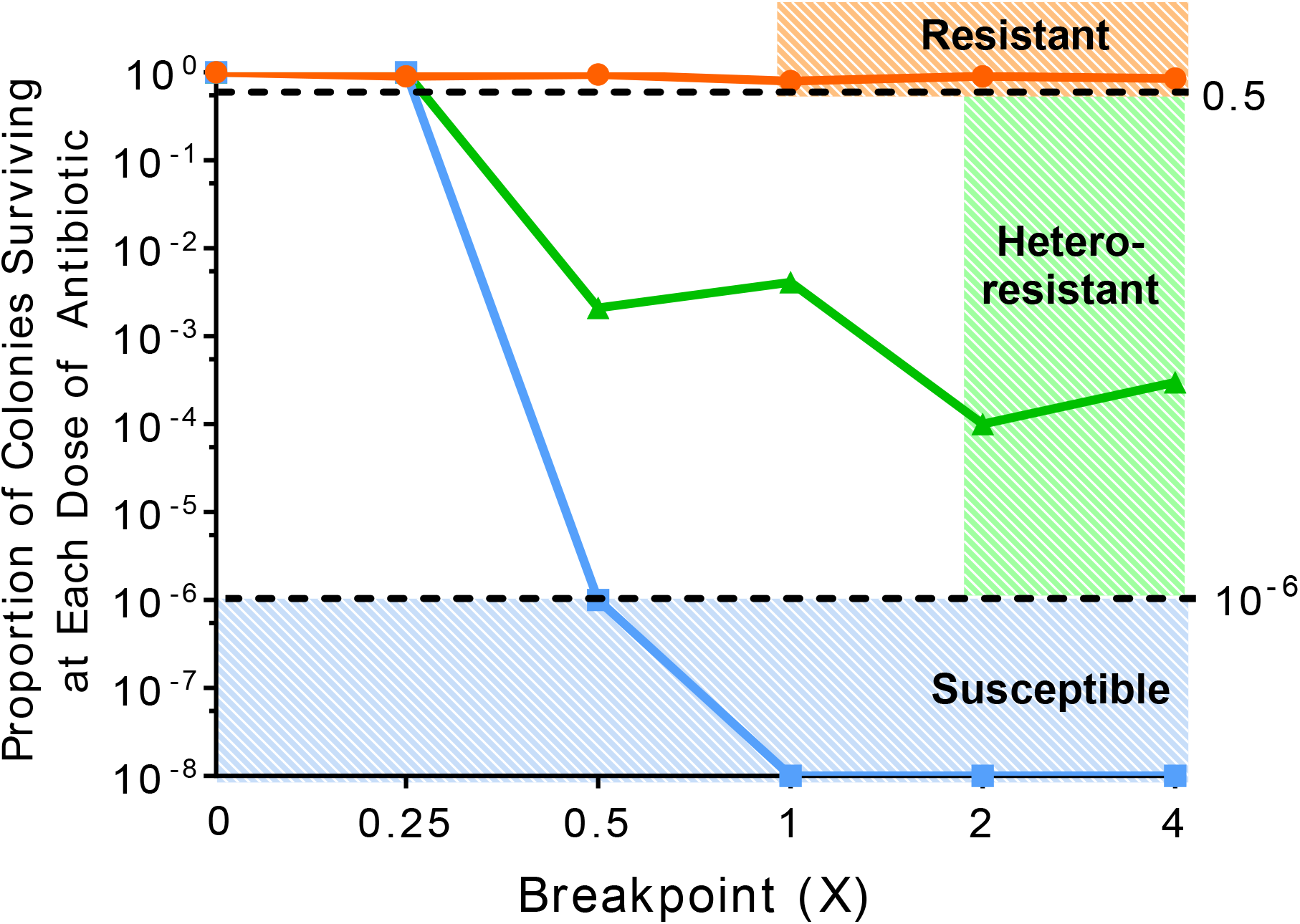
Designation of study strains by population analysis profile (PAP). A representative graph of a susceptible (blue), heteroresistant (green)and resistant (orange) isolate from this study. Isolates are designated resistant if at 1X or above the breakpoint there is survival of at least 50% (0.5) of the population. If the isolate is not resistant, it will be designated heteroresistant if at least 1 in 10^-6^ bacteria survive at 2X or above the breakpoint. If the isolate is neither resistant or heteroresistant, it will fall below the 10^-6^ at or before the breakpoint and is designated susceptible.

